# DeepPSC (protein structure camera): computer vision-based protein backbone structure reconstruction from alpha carbon trace as a case study

**DOI:** 10.1101/2020.08.12.247312

**Authors:** Xing Zhang, Junwen Luo, Yi Cai, Wei Zhu, Xiaofeng Yang, Hongmin Cai, Zhanglin Lin

## Abstract

Deep learning has been increasingly used in protein tertiary structure prediction, a major goal in life science. However, all the algorithms developed so far mostly use protein sequences as input, whereas the vast amount of protein tertiary structure information available in the Protein Data Bank (PDB) database remains largely unused, because of the inherent complexity of 3D data computation. In this study, we propose Protein Structure Camera (PSC) as an approach to convert protein structures into images. As a case study, we developed a deep learning method incorporating PSC (DeepPSC) to reconstruct protein backbone structures from alpha carbon traces. DeepPSC outperformed all the methods currently available for this task. This PSC approach provides a useful tool for protein structure representation, and for the application of deep learning in protein structure prediction and protein engineering.

## Introduction

Protein structure determination is an ongoing issue and a major goal in life science that has captivated the attention of scientists for decades. Experimentally, protein structures have been mostly determined by X-ray diffraction crystallography^1^, and to a less extent by nuclear magnetic resonance spectroscopy^2^. In recent years, cryo-electron microscopy (EM) has also been increasingly used for structure determination^3^. As an alternative to experimental methods, computational methods have also been developed for predicting protein structures from protein sequences, and deep learning has recently been applied to this prediction problem^4-6^. In particular, DeepMind proposed a method called AlphaFold^4^, which significantly outperformed all previous prediction methods. A number of algorithms that extract features from protein primary sequences for the purpose of protein function prediction and protein engineering, *e*.*g*., UniRep^7^ and TAPE^8^, represent a further advancement in the field. Other applications of deep learning include protein fold recognition^9^, and the predictions of protein secondary structures^10^, protein functions^11^, and drug protein interactions^12^.

However, all the deep learning methods developed so far utilize only protein sequences as input, whereas the vast amount of protein tertiary structure information available in the rapidly expanding PDB database has not been sufficiently exploited in the calculations, due to its complexity. There are presently three common coarse approximation approaches for protein structure representation, namely, *k* nearest residues^13^, distance or contact maps^14^, and 3D grids^15^, but all with limited utility. Thus, we are interested in the following question: how to utilize protein structure information in deep learning?

It is well known that images can be efficiently processed by deep learning, and particularly in recent years, convolutional neural networks (CNN) have been successfully used in an array of computer vision tasks such as image classification^16^, object detection^17^, and face recognition^18^. CNN can understand an object in the Euclidean space by extracting visual features from the corresponding image^19,20^. Fig. 1a shows a typical workflow of computer vision-based image classification. Here we propose a “Protein Structure Camera” (PSC) approach for converting protein tertiary structures into images for computer vision processing. In PSC, we used a 16 Å × 16 Å × 16 Å sliding cubic window centered on the alpha carbons of the amino acid residues (Cα) to dissect a protein structure (Fig. 1b). This was then turned into a group of compressed two dimensional 16 Å× 16 Å images with a -8 Å to 8 Å depth range, which were then fed into a CNN and implemented into a deep learning-based network architecture, or DeepPSC (Fig. 1c).

**Figure 1.**
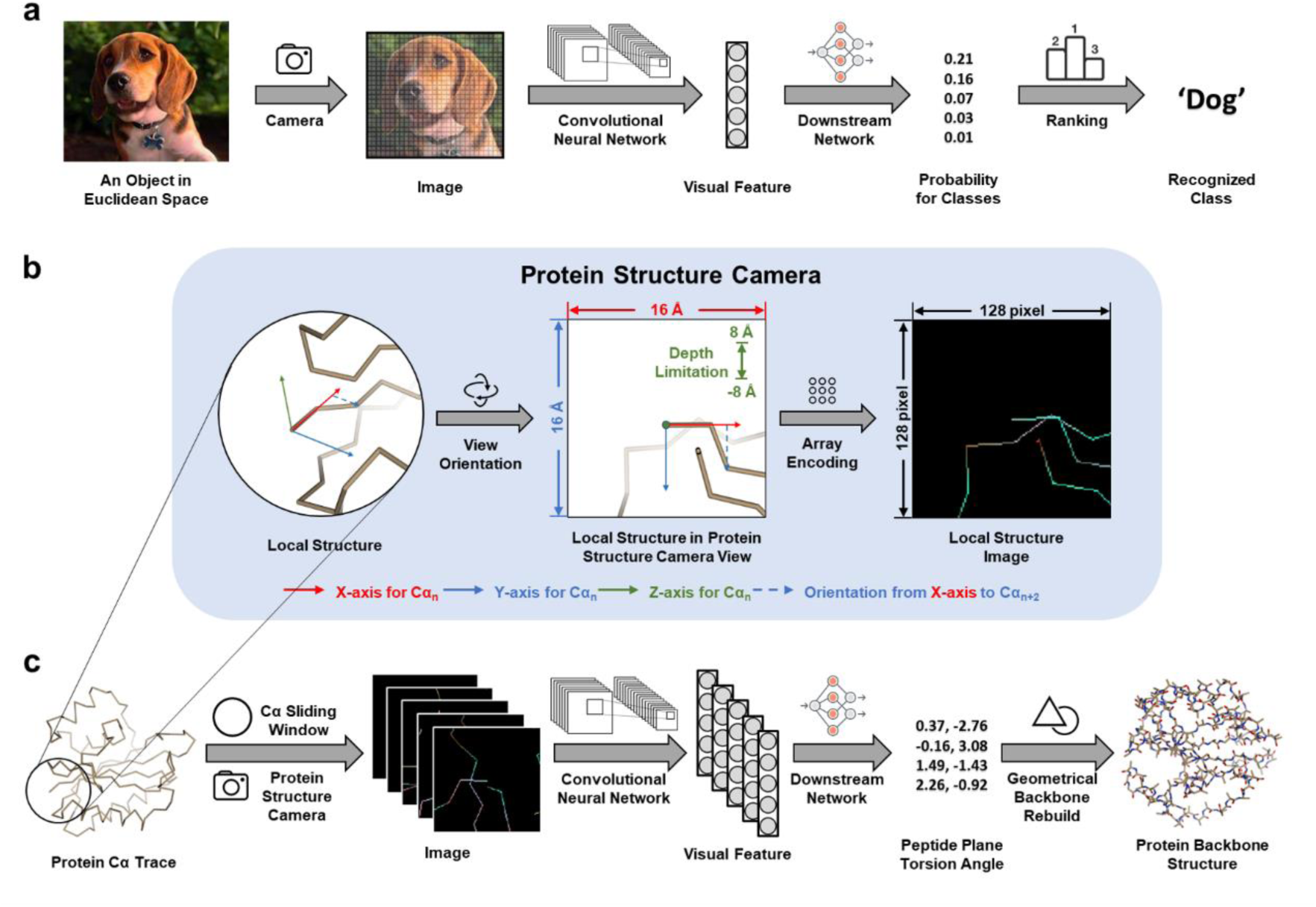
Schematic for the deep learning algorithm used in this work, or DeepPSC. a) The workflow of a typical computer vision task. b) Visualization of Protein Structure Camera (PSC) workflow. This figure is generated with the Chimera software ^50^. c) The workflow of the deep learning-based algorithm DeepPSC used in this work for reconstructing backbone structures from alpha-carbon traces.

As a case study, we applied this DeepPSC for reconstructing protein backbone structures (containing atoms C, N, O, Cβ in addition to Cα) based on Cα traces, which is an important task for protein structure determination by experimental means and for protein structure prediction by computational approaches. Several protein structure refinement methods have been developed for the analysis of EM images to generate high-quality structure models, such as PHENIX^21^ and Coot^22^. Within these algorithms, the positions of the Cα, which are the atoms that can be located with the highest accuracy, are determined first. Subsequently the backbone structure and then the full atom model are generated. Similarly, many computational algorithms predict the Cα trace as a preliminary reduced model. PHENIX ensembles PULCHRA^23^ for backbone reconstruction, which uses a simple force field and steepest descent minimization. Coot ensembles CALPHA^24,25^, which is based on a library of backbone fragments compiled from experimentally determined structures. The widely used computational protein structure prediction platform I-TASSER^26^ ensembles REMO^27^, which directly reconstructs full-atom models (including the backbones) from a backbone isomer library. Similar library-based methods include BBQ^28^, SABBAC^29^, and PD2^30^, which often achieve better performance than PULCHRA or REMO. These three backbone structure reconstruction methods have also been applied for experimental structure determination^31^, although they have not been incorporated in PHENIX or Coot. A significant limitation of the library-based methods, however, is that the wide range of conformations of protein backbones cannot be sufficiently represented by the limited number of fragments in the libraries.

In this work, we found that our DeepPSC approach outperformed all the previously reported methods for backbone reconstruction, including the benchmark PD2, and the ablation tests showed that the visual feature extracted from the protein structure images provided the main contribution for the improved performance.

## Results

### Represent Cα trace as images by protein structure camera

The PSC concept is shown in Fig. 1b. Given an Cα-trace {*Cα*_1_, *Cα*_2_, …, *Cα*_*L*_}, where *Cα*_*n*_ ∈ ℝ^3^ is the coordinate of the *n*^*th*^ Cα atom and *L* is the number of residues, PSC represents it as *L* images. Any given structural segment having *Cα*_*n*_ as the center requires a preset orientation and scale. We defined the orientation from *Cα*_*n*_ to *Cα*_*n*+1_ as the X-axis. The Y-axis was then determined by the orientation from the X-axis to *Cα*_*n*+2_, and the Z-axis was defined such as to build a left-hand Cartesian coordinate on the given local structural segment. We set the orientation directed from the positive to the negative regions of the Z-axis as the PSC view so that the orientations of different local structural segments could be normalized. For *Cα*_*L*−1_, the position on the Y-axis was determined by the orientation from the X-axis to *Cα*_*L*−2_. Similarly, for the last Cα, we defined the orientation from *Cα*_*L*_ to *Cα*_*L*−1_ as the X-axis and the orientation from the X-axis to *Cα*_*L*−2_ as the Y-axis. An enlarged view of Fig. 1b is given as Supplementary Fig. 1.

Since 8 Å is generally regarded as the interaction distance cutoff between two residues^32^, we used a sliding cubic window with a side length of 16 Å centered on the Cα, and a depth ranging from -8 Å to 8 Å was applied to the PSC view. Each PSC view was then encoded as an image with five channels, representing the Z-axis depth, the relative sequence position, and three key amino acid properties including hydrophobicity^33^, bulkiness^34^ and flexibility^35^, respectively. The resolution of the image is 128 × 128 pixels. In the image, each Cα was first encoded as a pixel, and a straight line was used to connect adjacent Cα pixels. The values of the properties along the straight line were interpolated from the two Cα pixel values. A given protein Cα trace was thus converted into a group of local structural images.

### Present protein backbone structure as peptide plane torsion angles

In this study, we represented the structure of the protein backbone as peptide plane torsions, as reported in a previous study^36^. For convenience, we denoted the C atom and N atom in the *n*^*th*^ of all *L* − 1 peptide planes by *C*_*n*_ ∈ ℝ^3^ and *N*_*n*_ ∈ ℝ^3^, respectively. Note that *N*_*n*_, the N atom in the *n*^*th*^ peptide, is actually the (*n* + 1)^*th*^ N atom of the protein backbone. Besides, we defined the vectors from atom *A* to atom *B* as 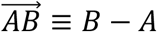, with the corresponding unit vector being 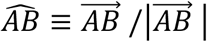. We applied a constraint that assumed *Cα*_*n*_, *C*_*n*_, *O*_*n*_, *N*_*n*_, *Cα*_*n*+1_ forming a standard peptide plane (*trans* or *cis*). Within the given *n*^*th*^ *trans* peptide plane, as shown in Fig.2a, the locations of *C*_*n*_, *O*_*n*_, *N*_*n*_ on the plane can be determined with a group of fixed lengths. For example, since |*Cα*_*n*_*PC*_*n*_| and |*PC*_*n*_*C*_*n*_| are fixed, we could locate *C*_*n*_ on the plane. Next, as shown in Fig. 2b, we used 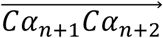 as a reference orientation to determine the torsion angles for each *n*^*th*^peptide plane, *i*.*e*., the torsion angle from 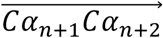 to 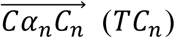, and that from 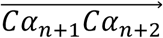 to 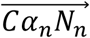 (*TN*_*n*_), where *TN*_*n*_ is approximately 180 degrees larger than *TC*_*n*_. The locations of *Cα*_*n*_, *C*_*n*_, *O*_*n*_, *N*_*n*_, *Cα*_*n*+1_ were determined from the combination of all this information. In the case of *cis* peptides, the fixed lengths are different from those of *trans* peptides, and *TN*_*n*_ is close to *TC*_*n*_. In all cases, *TO*_*n*_ is very close to *TC*_*n*_, therefore *TC*_*n*_ was used as an approximation of *TO*_*n*_ in all calculations. *TC*_*n*_ and *TN*_*n*_ were encoded in the form of sine and cosine in the final representations.

**Figure 2.**
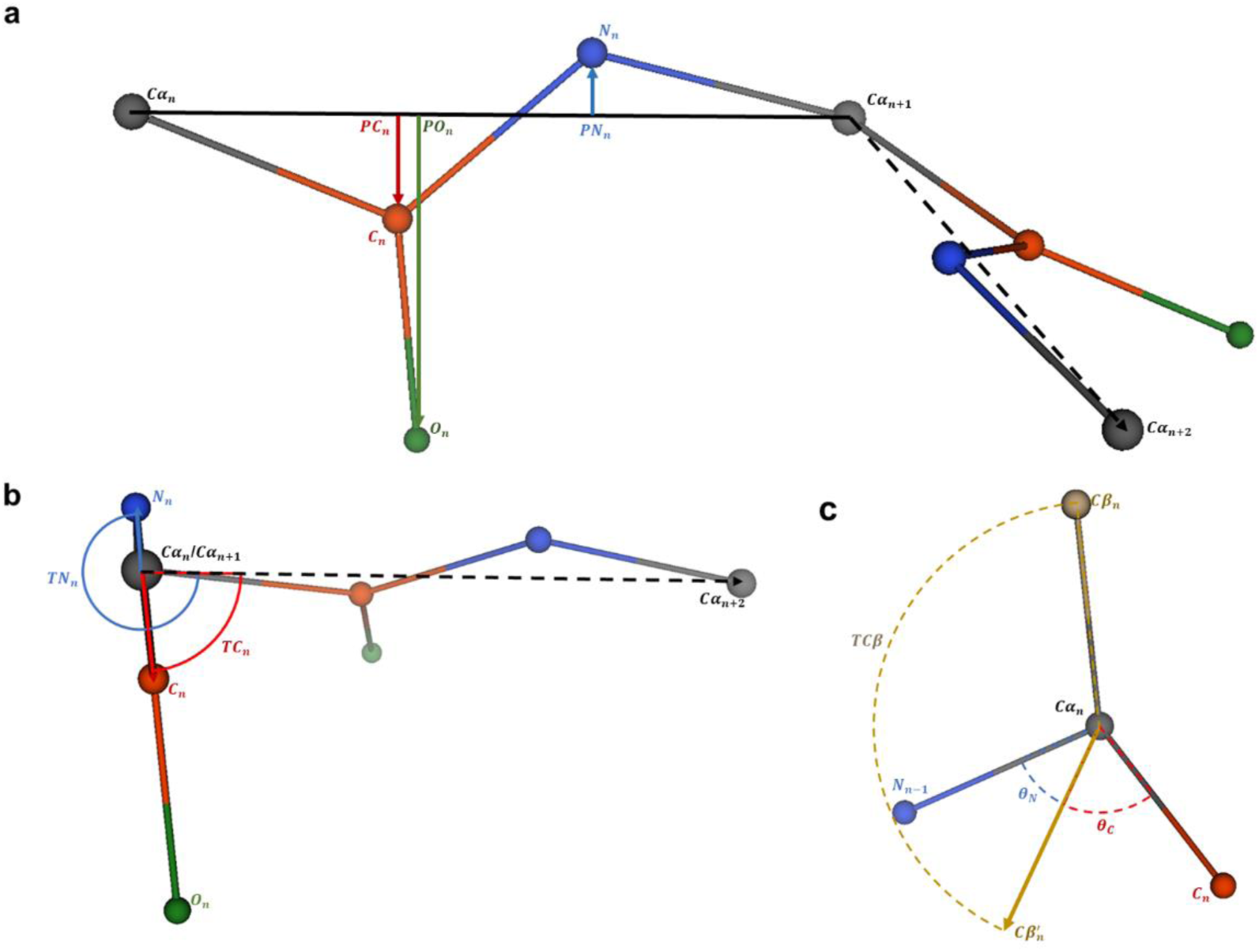
Diagrams of the backbone structure representation and rebuilding (only the case of a trans peptide plane is shown). a) Typical peptide plane conformation. *Cα*_*n*_, *C*_*n*_, *O*_*n*_, *N*_*n*_, *Cα*_*n*+1_ located on the *n*^*th*^ peptide plane of the protein structure. *PC*_*n*_, *PN*_*n*_, and *PO*_*n*_ are the projections of *C*_*n*_, *N*_*n*_, and *O*_*n*_ on 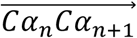, respectively. b) Side view from *Cα*_*n*_ to *C*_*n*+1_, in which *TC*_*n*_ and *TN*_*n*_ are the torsion angles from *Cα*_*n*+2_ to *C*_*n*_, and from *Cα*_*n*+2_ to *N*_*n*_, respectively, with 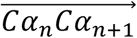 as the axis. c) Rebuilding process for *Cβ*_*n*_, using the constraint *θ*_*N*_ = *θ*_*C*_, fixed bond length 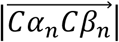, and fixed torsion angle *TCβ*.

Since the residues in proteins are L-amino acids, the coordinates of *Cβ*_*n*_ can be determined when *N*_*n*−1_, *Cα*_*n*_, and *C* are known. Fig. 2c shows the rebuilding process used in this work. We first set the position of the projection of 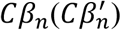 along the direction of the bisector of ∠*N*_*n*−1_*Cα*_*n*_*C*_*n*_, with a fixed bond length |*Cα*_*n*_*Cβ*_*n*_|. By rotating 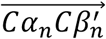 with the fixed angle *TCβ*, the location of *Cβ*_n_ was determined. Without constraints from the peptide plane, the first N atom and the last C and O atoms of a protein backbone are usually highly flexible, therefore our method could not predict the positions of these atoms.

Specific geometrical calculations for all the above representations and rebuilding are provided in Methods.

### Develop a deep neural network for PSC images

As shown in Fig. 3, the deep neural network implemented in DeepPSC takes local structure images as the input, and calculate the peptide plane torsions as the output. We first adopted ResNet50^37^, the most used convolutional neural network for computer vision processing, to extract visual features from the images (Supplementary Fig. 2), which are labelled as “local structural features”. Then, we used a bidirectional long short-term memory module (Bi-LSTM)^38,39^, the most used recurrent neural network module for sequence modelling, to sequentially pass information between the extracted local structural features (Supplementary Fig. 3a). The outputs of this module were expected to mainly represent the local structures but they also contain sequential context information, and are labeled as “globalized local structural features”. Afterward, considering that a single peptide plane is constructed by two adjacent residues, we paired every globalized local structural feature with the next one in the amino acid sequence as the “peptide plane feature”. Finally, we used a multilayer perceptron (MLP)^40^, a typical neural network module, to predict peptide plane torsions from the peptide plane features (Supplementary Fig. 3b).

**Figure 3.**
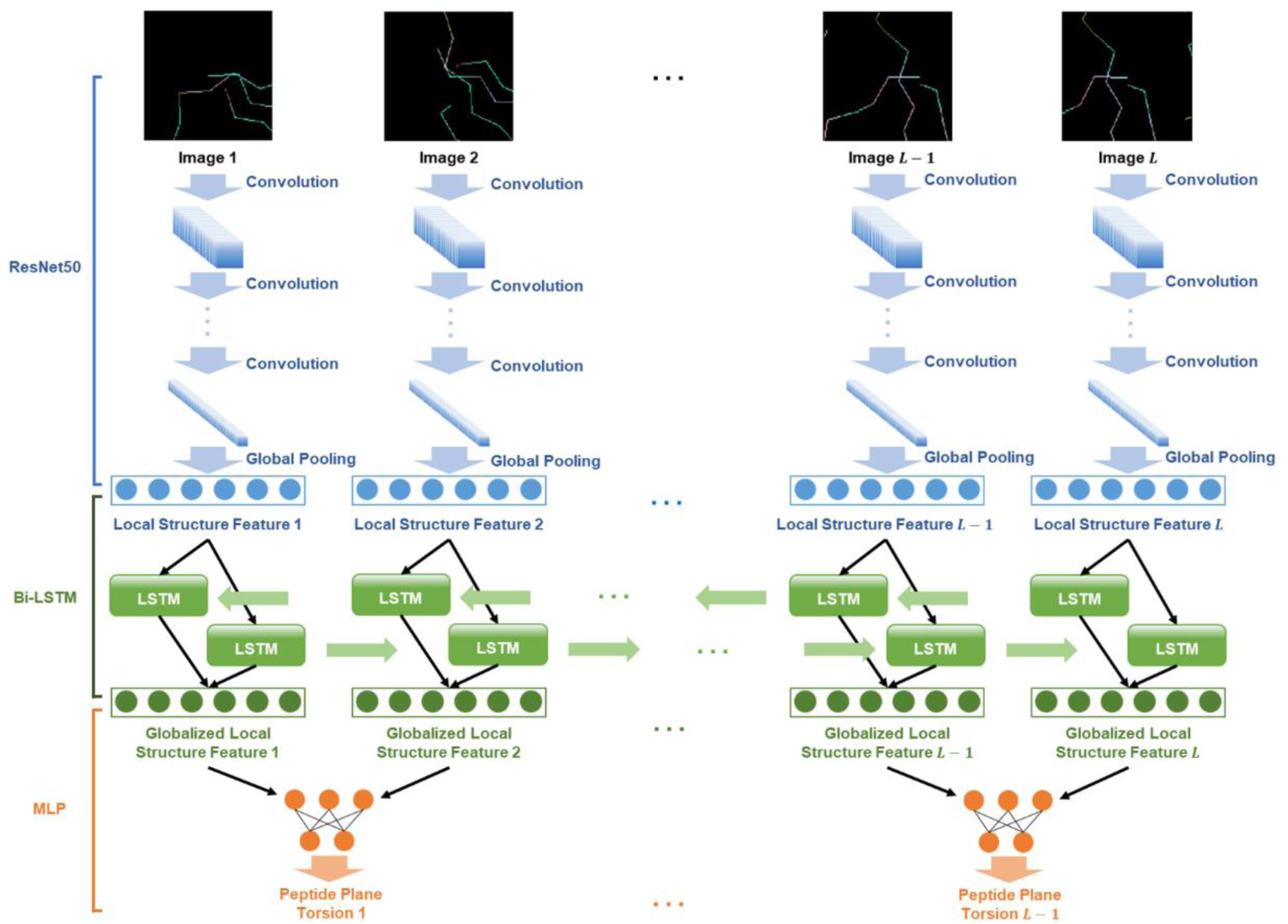
Network Architecture of DeepPSC. ResNet50 was used to extract visual features from images as local structure features. All local structure features were then fed into a Bi-LSTM module for information globalization among residues, yielding globalized local structure features. Finally, an MLP module was used to predict peptide plane torsions for pairs of the above adjacent globalized local structure features.

To compare our DeepPSC method with previously reported protein structure representation methods, we additionally built two baseline methods. In the first baseline, we used the k nearest residues method^13^ to represent the Cα trace, and to encode the network input. To maintain the input information as close to that of our method as possible, we enriched the representation by adding relative protein positions and residue properties. For this baseline, an MLP module (Supplementary Fig. 4) was used instead of ResNet50, to extract local structural features for the input format, since the latter cannot process this baseline input^13^. For the second baseline, protein structures were represented as distance maps and processed with a CNN (Supplementary Fig. 5), as previously reported, without any modifications^14^.

### DeepPSC outperforms other standard backbone reconstruction methods

We performed the 10-fold cross validation process on the three network architectures (DeepPSC, and the two baselines), and obtained 10 models for each architecture, for a total of 30 models. Next, we applied each of these models to the test set and obtained the predicted torsion angles as outputs. Subsequently, for each architecture, we took the average of the outputs of the 10 cross validation models as the “ensemble model”. Then the outputs of the ten models and that of the ensemble model were used to rebuild the backbone structures together with the corresponding Cα traces, and these rebuilt models were evaluated with the three performance criteria, RMSD_100_, GDT_P0.2, and RAMA outliers (Table 1). The average performance of the ten models for each architecture was calculated and shown as the “single model” performance, with the standard deviation of the single model performance indicating the robustness of the network architecture. Finally, we compared the performance of these architectures to that of PD2, BBQ, SABBAC and PULCHRA (Table 1).

**Table 1.** Overview of the results for various backbone reconstruction methods.

| Methods | Models | Mean<br>RMSD<sub>100</sub> (Å) | Mean<br>GDT_P0.2 (%) | Mean RAMA<br>outliers (%) |
| --- | --- | --- | --- | --- |
| Rebuilt from<br>PDB | — | 0.040 | 95.76 | 0.22 |
| DeepPSC | Ensemble<br>model | **0.076** | **88.18** | **0.23** |
|  | Single model | 0.079±0.001 | 87.87±0.08 | 0.22±0.03 |
| Baseline 1<br>(*k* nearest<br>residues) | Ensemble<br>model | 0.101 | 83.44 | 0.50 |
|  | Single model | 0.108±0.001 | 82.16±0.25 | 0.71±0.07 |
| Baseline 2<br>(distance map) | Ensemble<br>model | 0.289 | 54.58 | 5.23 |
|  | Single model | 0.289±0.001 | 54.42±0.48 | 5.30±0.17 |
| PD2 | — | 0.149 | 73.26 | 0.90 |
| BBQ | — | 0.156 | 71.82 | 3.03 |
| SABBAC* | — | 0.201 | 58.30 | 1.94 |
| PULCHRA | — | 0.221 | 52.15 | 2.20 |

\*SABBAC failed to process one of the 21 structures in the test set. The results shown here were obtained with the other 20 test structures.

Generally, protein structures with resolution smaller than 2.0 Å are regarded as high-quality structures^30^. According to the official statistics of PDB, up to July 27, 2020, the median resolution of X-ray crystallography structures in the database is 2.03 Å. For a typical 2.0 Å crystallographic model, the average error on atomic coordinates is lower than 0.2 Å^41^. Therefore, we considered 0.2 Å as the benchmark in our performance evaluation. Accordingly, we set the GDT cutoff at 0.2 Å to calculate the percentage of atoms that can be regarded as acceptable in a high-quality structure.

Based on the mean RMSD_100_, and the GDT_P0.2 and RAMA outliers percentages as shown in Table 1, the backbone structures predicted by the ensemble model obtained with DeepPSC clearly outperformed those predicted by the baseline methods as well as the various traditional methods (PD2, BBQ, SABBAC and PULCHRA), in all three criteria. In particular, the performance of baseline 1, which was devoid of the image features of DeepPSC, suggested that the visual features extracted in DeepPSC were the main factor for its improved performance. By comparing the results for the Rebuilt model (directly from the PDB) and the ensemble model of DeepPSC, it could be deduced that the deviations observed in DeepPSC consisted of two elements: (i) the first was represented by the deviations introduced during the rebuilding process per se, which were the deviations between the ideal peptide plane conformations and the experimentally determined peptide plane conformations; (ii) the second is represented by the deviations induced by the model fitting in DeepPSC. Therefore, future developments should focus on devising an alternative strategy in lieu of the peptide plane assumption.

The distributions of the atomic coordinate deviations were also used to calculate the GDT_P0.2 scores of DeepPSC, PD2, PULCHRA and the two baselines (Fig. 4), which clearly show that the backbones reconstructed by DeepPSC were more accurate than those obtained with PD2, PULCHRA and the two baselines. Lastly, the Ramachandran plots of the reconstructions obtained with different methods clearly showed that the backbone structures reconstructed by DeepPSC were the most reasonable among all methods (Fig. 5). In particular, none of the glycine and proline residues in the backbones obtained from DeepPSC were classified as outliers, consistent with the experimentally determined PDB structures, whereas many of these residues resulted as outliers in the backbones obtained by PD2 and PULCHRA, as well as by baseline 2. It is noteworthy that for baseline 2, the dihedral angles share a “S-shape” distribution pattern for all the four types of residues (general, glycine, pre-proline, and proline), which is consistent with the poor network fitting of this type of protein structure representation, as shown in Supplementary Fig. 6.

**Figure 4.**
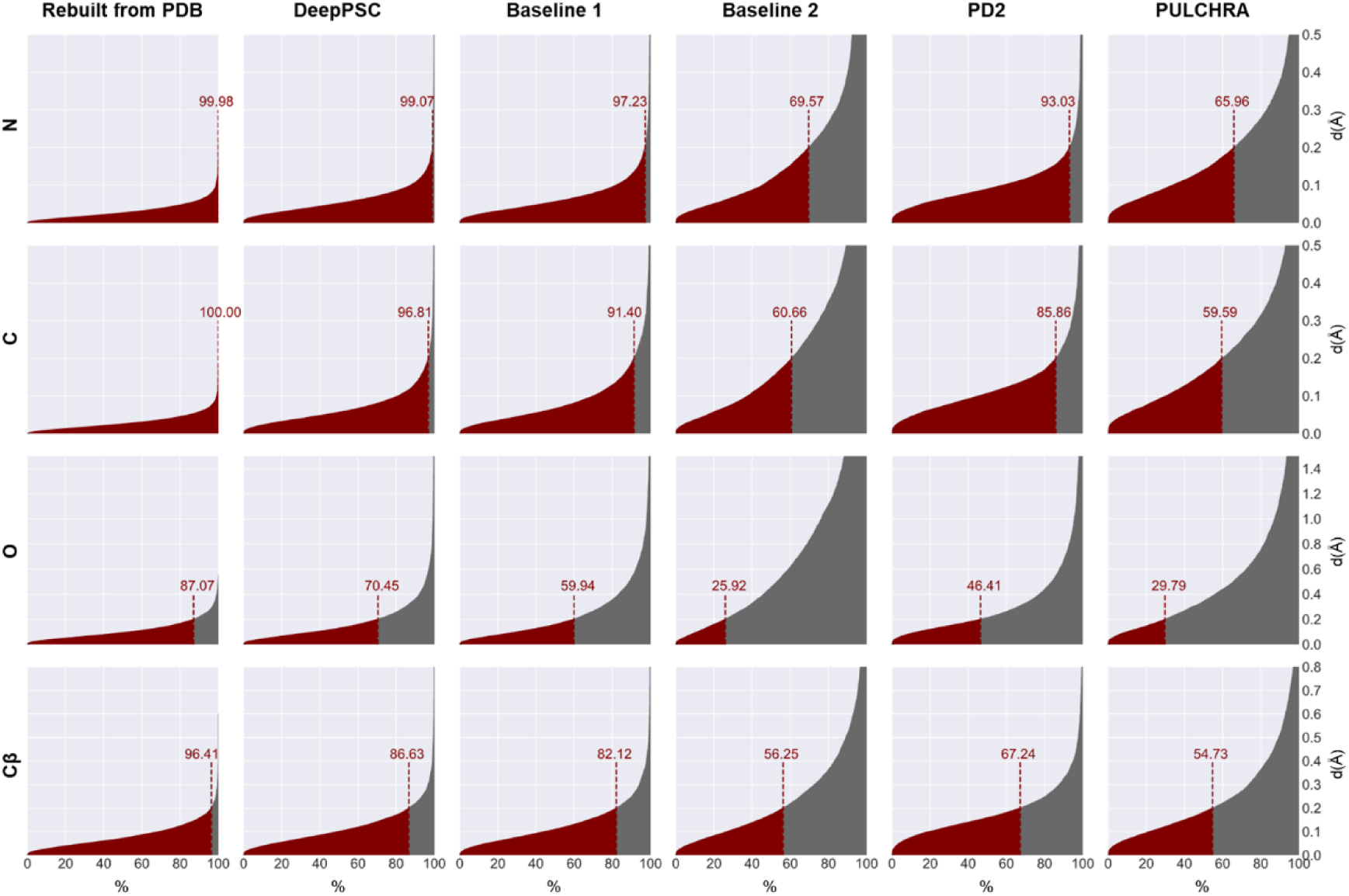
Distributions of the atomic coordinate deviations (rows) of the various reconstruction methods (columns). The GDT scores for 0.2 Å cutoff are indicated in the plots.

**Figure 5.**
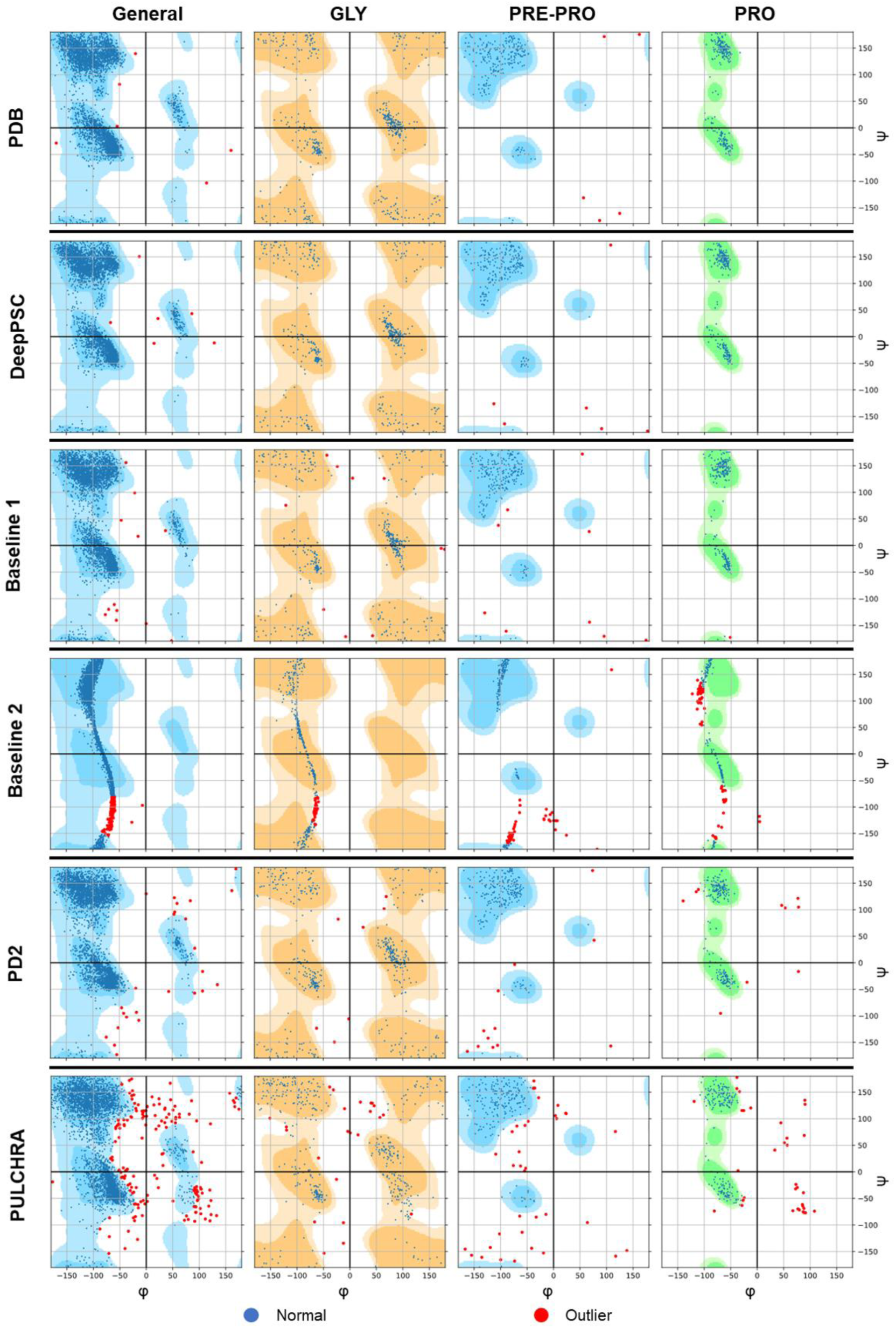
Ramachandran plots of the reconstructions obtained with different methods compared to the original structures (PDB). Rows represent the different methods and columns represent all residues (General), glycines (Gly), the residues preceding prolines (Pre-Pro), and prolines (Pro). By taking the reference distributions in the backgrounds, residues are classified as normal residues (blue) or outliers (red).

## Conclusions

We consider protein structure representation as a critical problem in applying deep learning for reliable protein structure prediction, and for related endeavors such as protein design. Our protein structure camera (PSC) approach provides a step forward in protein structure representations, and toward enabling more sophisticated applications of deep learning in biology.

## Methods

### Geometrical calculation

In this study, we represented C atoms and N atoms in peptide planes as torsion angles by:

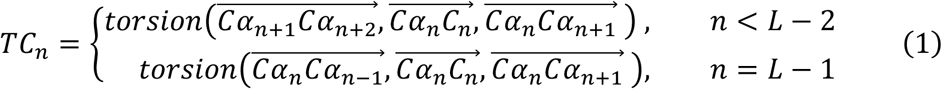

and

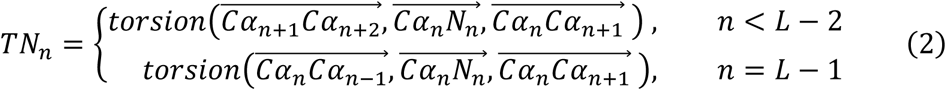

in which the torsion angle from 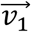 to 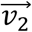 with 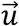 as axis was calculated by:

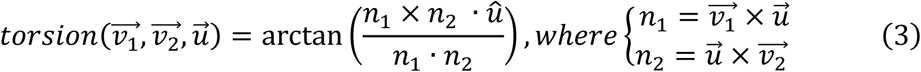

In the rebuilding process, the orientation of *C*_*n*_ and *N*_*n*_ was determined by rotating the *n*^*th*^ peptide plane with 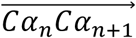 as the axis:

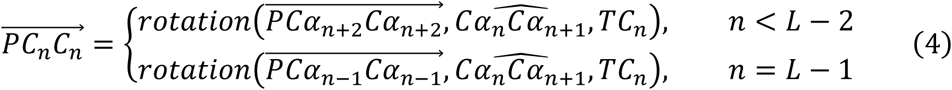

and

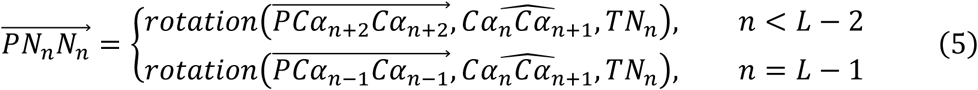

where *PCα*_*n*+2_ and *PCα*_*n*−1_ is the projections of *Cα*_*n*+2_ and *Cα*_*n*−1_ on 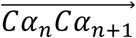, respectively. The rotation was calculated by Rodrigues-Gibbs Formulation^42^:

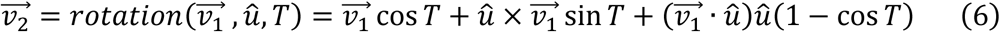

in which 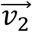 was obtained by rotate 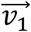 with a unit vector û as axis and *T* as torsion angle. We assumed 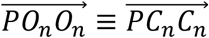 using the ideal peptide plane conformation. Afterward the relative locations from atoms *C*_*n*_, *N*_*n*_, and *O*_*n*_ to *Cα*_*n*_ were respectively determined by:

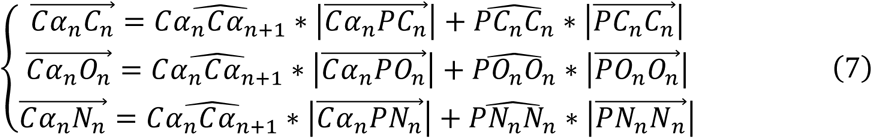

where 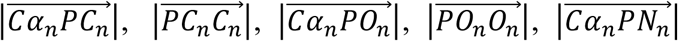, and 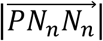 are a group of fixed length estimated from training data (listed in Table S1). Note that there are two type of peptide planes and the fixed lengths are correspondingly different. In the *trans* peptide plane, the distance between adjacent Cα is approximately 3.8 Å while for the *cis* peptide plane it is approximately 3.0 Å. Therefore, we rebuilt the *n*^*th*^ peptide plane with fixed lengths for the *trans* peptide plane when 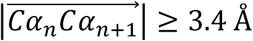, otherwise with fixed lengths for *cis* peptide plane.

Finally, the coordinates were determined as:

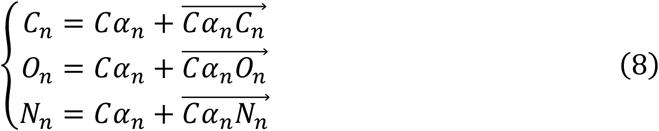

The rebuilt atoms *C*_*n*_, *N*_*n*−1_ and known *Cα*_*n*_ in a residue were used to rebuild atom *Cβ*_*n*_ ∈ ℝ^3^ with the following process. First, we initialized 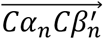 at the middle between 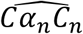 and 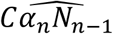 by:

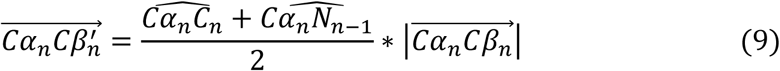

Afterward, 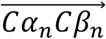 was determined by rotating 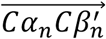 with 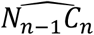 as axis and *TCβ* as torsion angle:

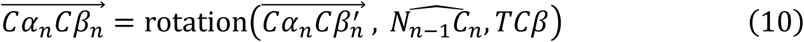

We then calculated *Cβ*_*n*_ by:

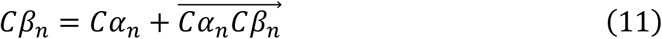

The fixed length 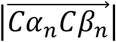 and fixed angle *TCβ* used above are constants estimated from the training data (listed in Supplementary Table 1).

#### Dataset

We selected a subgroup from the protein structures reported in the PDB, to build a tertiary structure dataset. A PDB entry was not included in this subgroup if: (i) the structure was not determined by X-ray crystallography; (ii) the entry has a number of residues fewer than 15 or higher than 800; (iii) the entry has missing atoms in the backbone or unnatural residues; (iv) the entry has sequence identity higher than 40% with another entry included in the subgroup^30^. This resulted in the construction of a non-redundant dataset containing 10,302 protein structures.

#### Model training

We used 10-fold cross validation in training our models. The whole dataset was randomly and equally separated into ten sub-datasets. We routinely used one sub-dataset as the validation set and all the other nine sub-datasets as the model training sets. All the models were trained for 30 epochs using mean square error (MSE) as the loss function and the Adam optimizer^43^. The training batch size for the local structure embedding block was the total number of residues of the input structures, thus it was dynamic even if the number of input structures was fixed. To maintain the training batch relatively stable, we split all training structures as batches containing 1∼5 structures with approximately 800 residues. The learning rate was set to 0.0003 for the first 3 epochs, and then was adjusted according to the cosine-annealing schedule^44^ in the following epochs. The trained models were validated with the corresponding validation set after every epoch. The curves of validation loss showed that all the models of DeepPSC and baseline 1 steadily converged at the 30^th^ epoch, while the models of baseline 2 showed poor fitting (Supplementary Fig. 6). The network construction and model training were implemented with PyTorch, an open source machine learning framework. All the details we have not mention follow the default setting of PyTorch.

#### Performance criteria

In this study the performance of various methods was evaluated on the basis of three criteria, *e*.*g*., root mean square deviation (RMSD)^45^, global distance test (GDT)^46^, and Ramachandran (RAMA) outliers^47^. RMSD is one of the most used criteria to measure the similarity between two structures ^45^, and is calculated as follows:

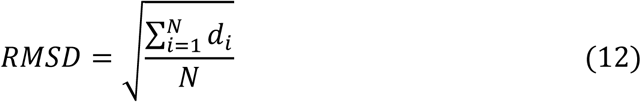

where *d*_*i*_ is the coordinate deviation between atom *i* in two structures, and *N* is the total number of atoms. Considering that RMSD usually increases as the number of atoms of a protein increases^48^, this value is usually normalized as RMSD_100_, which describes the same deviation in 100 atoms^49^, and is calculated as follows:

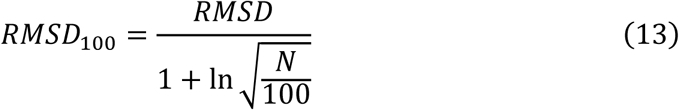

where *N* is the number of atoms.

RMSD is, however, strongly affected by the parts in the structures that deviate the most, therefore it often fails to represent the deviations of most of the atoms. Aimed at alleviating this problem, a community-wide experiment called CASP (critical assessment of techniques for Protein Structure Prediction)^46^ have been using a different indicator, the Global Distance Test (GDT), as their main assessment method for ranking protein structure prediction methods. GDT scores are calculated as the percentage of atoms that have distance deviations smaller than the preset distance cutoffs. Cutoffs for GDT in CASP is usually set to 1, 2, 4, and 8 Å. In this study, the cutoff was set to 0.2 Å, and the GDT was labeled as GDT_P0.2.

The Ramachandran Plot is a statistical reference distribution of the combination of the backbone dihedral angles in proteins^47^. In a Ramachandran Plot, one can classify residues in a given protein backbone structure as ‘core’, ‘allowed’, and ‘outliers’. The percentage of outliers (RAMA outliers) is used to assess protein backbone structure uncertainty.

## Acknowledgements

This work was supported by the National Key R&D Program of China (2018YFA0901000), and the Guangzhou Science and Technology Program key projects (201904020016). Dr. Hongmin Cai acknowledges the support by the Key-Area Research and Development of Guangdong Province under Grant (2020B010166002, 2020B1111190001), the National Natural Science Foundation of China (61771007), the Health & Medical Collaborative Innovation Project of Guangzhou City (201803010021, 202002020049).

## Author contributions

X.Z. contributed to the experimental design, methodology, coding, data analysis and writing of the original draft. J.L. contributed to the methodology, coding and data analysis. Y.C contributed to the data analysis. X.Y. contributed to the data analysis and writing (review and revision). W.Z. contributed to the data analysis. H.C. contributed to the methodology, data analysis and writing (review and revision), and was partially responsible for supervision. Z.L. was responsible for the experimental design, supervision as well as funding acquisition, and contributed to the data analysis, writing (review and revision). All authors reviewed and approved the final manuscript.

## Competing interests

The authors declare no competing interests.

## Code and data availability

All source codes and models of DeepPSC are openly available on GitHub (https://github.com/EricZhangSCUT/DeepPSC), together with the PDB ID lists of all involved datasets.

## Notes

### Competing Interest Statement

The authors have declared no competing interest.

### Summary of Updates

Geometrical calculation detail updated to clarify the definition of peptide plane torsion angles; Supplemental files updated.

## Reference

1 Smyth, M. S. & Martin, J. H. X-ray crystallography. Mol. Pathol. 53, 8–14 (2000).

2 Voula Kanelis, J. D.F.-K. & Lewis E. Kay. Multidimensional NMR Methods for Protein Structure Determination. IUBMB Life 52, 291–302 (2001).

3 Cheng, Y. Single-particle cryo-EM - How did it get here and where will it go. Science 361, 876–880 (2018).

4 Senior, A. W. et al. Improved protein structure prediction using potentials from deep learning. Nature 577, 706–710 (2020).

5 AlQuraishi, M. End-to-end differentiable learning of protein structure. Cell Syst. 8, 292–301 e293 (2019).

6 Wang, S., Sun, S., Li, Z., Zhang, R. & Xu, J. Accurate de novo prediction of protein contact map by ultra-deep learning model. PLoS Comput. Biol. 13, e1005324 (2017).

7 Alley, E. C., Khimulya, G., Biswas, S., AlQuraishi, M. & Church, G. M. Unified rational protein engineering with sequence-based deep representation learning. Nat. Methods 16, 1315–1322 (2019).

8 Rao R, B.N., Thomas N, et al. Evaluating protein transfer learning with TAPE. Preprint at https://arxiv.org/abs/1906.08230 (2019).

9 Hou, J., Adhikari, B. & Cheng, J. DeepSF: deep convolutional neural network for mapping protein sequences to folds. Bioinformatics 34, 1295–1303 (2018).

10 Wang, S., Peng, J., Ma, J. & Xu, J. Protein secondary structure prediction using deep convolutional neural fields. Sci. Rep. 6, 18962 (2016).

11 Kulmanov, M., Khan, M. A., Hoehndorf, R. & Wren, J. DeepGO: predicting protein functions from sequence and interactions using a deep ontology-aware classifier. Bioinformatics 34, 660–668 (2018).

12 Tsubaki, M., Tomii, K. & Sese, J. Compound-protein interaction prediction with end-to-end learning of neural networks for graphs and sequences. Bioinformatics 35, 309–318 (2019).

13 Wang, J., Cao, H., Zhang, J. Z. H. & Qi, Y. Computational protein design with deep learning neural networks. Sci. Rep. 8, 6349 (2018).

14 Zheng, S., Li, Y., Chen, S., Xu, J. & Yang, Y. Predicting drug–protein interaction using quasi-visual question answering system. Nat. Mach. Intell. 2, 134–140 (2020).

15 Stepniewska-Dziubinska, M. M., Zielenkiewicz, P. & Siedlecki, P. Development and evaluation of a deep learning model for protein-ligand binding affinity prediction. Bioinformatics 34, 3666–3674 (2018).

16 Krizhevsky, A., Sutskever, I. & Hinton, G. E. Imagenet classification with deep convolutional neural networks. NeurIPS 1097-1105 (2012).

17 Zhao, Z.Q., Zheng, P., Xu, S.t. & Wu, X. Object detection with deep learning: A review. IEEE Trans. Neural Netw. Learn. Syst. 30, 3212–3232 (2019).

18 Sun, Y., Chen, Y., Wang, X. & Tang, X. Deep learning face representation by joint identification-verification. NeurIPS 1988-1996 (2014).

19 Mallat, S. Understanding deep convolutional networks. Philos. Trans. A Math. Phys. Eng. Sci. 374, 20150203 (2016).

20 Zeiler, M. D. & Fergus, R. Visualizing and understanding convolutional networks. European conference on computer vision 818–833 (2014).

21 Adams, P. D. et al. PHENIX: a comprehensive Python-based system for macromolecular structure solution. Acta Crystallogr. D Biol. Crystallogr. 66, 213–221 (2010).

22 Emsley, P. & Cowtan, K. Coot: model-building tools for molecular graphics. Acta Crystallogr. D Biol. Crystallogr. 60, 2126–2132 (2004).

23 Rotkiewicz, P. & Skolnick, J. Fast procedure for reconstruction of full-atom protein models from reduced representations. J. Comput. Chem. 29, 1460–1465 (2008).

24 Esnouf, R. M. Polyalanine Reconstruction from Ca Positions Using the Program CALPHA Can Aid Initial Phasing of Data by Molecular Replacement Procedures. Acta Crystallogr. D Biol. Crystallogr. 53, 665–672 (1997).

25 Emsley, P., Lohkamp, B., Scott, W. G. & Cowtan, K. Features and development of Coot. Acta Crystallogr. D Biol. Crystallogr. 66, 486–501 (2010).

26 Roy, A., Kucukural, A. & Zhang, Y. I-TASSER: a unified platform for automated protein structure and function prediction. Nat. Protoc. 5, 725–738 (2010).

27 Li, Y. & Zhang, Y. REMO: A new protocol to refine full atomic protein models from C-alpha traces by optimizing hydrogen-bonding networks. Proteins 76, 665–676 (2009).

28 Gront, D., Kmiecik, S. & Kolinski, A. Backbone building from quadrilaterals: a fast and accurate algorithm for protein backbone reconstruction from alpha carbon coordinates. J. Comput. Chem. 28, 1593–1597 (2007).

29 Maupetit, J., Gautier, R. & Tuffery, P. SABBAC: online Structural Alphabet-based protein BackBone reconstruction from Alpha-Carbon trace. Nucleic Acids Res. 34, W147–151 (2006).

30 Moore, B. L., Kelley, L. A., Barber, J., Murray, J. W. & MacDonald, J. T. High-quality protein backbone reconstruction from alpha carbons using Gaussian mixture models. J. Comput. Chem. 34, 1881–1889 (2013).

31 Zhang, R. et al. 4.4 A cryo-EM structure of an enveloped alphavirus Venezuelan equine encephalitis virus. EMBO J. 30, 3854–3863 (2011).

32 Xiong, D., Zeng, J. & Gong, H. A deep learning framework for improving long-range residue-residue contact prediction using a hierarchical strategy. Bioinformatics 33, 2675–2683 (2017).

33 Kyte, J. & Russell F. Doolittle. A simple method for displaying the hydropathic character of a protein. J. of Mol. Biol. 157, 105–132 (1982).

34 Zimmerman, J. M., Naomi Eliezer & R. Simha. The characterization of amino acid sequences in proteins by statistical methods. J. of Theor. Biol. 21.2, 170–201 (1968).

35 Huang, F. & Nau, W. M. A conformational flexibility scale for amino acids in peptides. Angew. Chem. Int. Ed. Engl. 42, 2269–2272 (2003).

36 Payne, P. W. Reconstruction of protein conformations from estimated positions of the Cα coordinates. Protein Sci. 2(3), 315–324 (1993).

37 He, K., Zhang, X., Ren, S. & Sun, J. Deep residual learning for image recognition. Proc. IEEE Comput. Soc. Conf. Comput. Vis. Pattern Recognit. 770–778 (2016).

38 Graves, A. & Schmidhuber, J. Framewise phoneme classification with bidirectional LSTM and other neural network architectures. Neural Netw. 18, 602–610 (2005).

39 Hochreiter, S. & Schmidhuber, J. Long short-term memory. Neural Comput. 9, 1735–1780 (1997).

40 Rumelhart, D. E., Hinton, G. E. & Williams, R. J. Parallel distributed processing: explorations in the microstructure of cognition, vol. 1 318–362 (MIT Press, Cambridge, 1986).

41 Scapin, G., Potter, C. S. & Carragher, B. Cryo-EM for small molecules discovery, design, understanding, and application. Cell Chem. Biol. 25, 1318–1325 (2018).

42 Parsons, J., Holmes, J. B., Rojas, J. M., Tsai, J. & Strauss, C. E. Practical conversion from torsion space to Cartesian space for in silico protein synthesis. J. Comput. Chem. 26, 1063–1068 (2005).

43 Kingma, D. P. & Jimmy Ba. Adam: A method for stochastic optimization. Preprint at https://arxiv.org/abs/1412.6980 (2014).

44 Loshchilov, I. & Frank Hutter. SGDR: Stochastic Gradient Descent with warm Restarts. Preprint at https://arxiv.org/abs/1608.03983 (2016).

45 Kufareva, I. & Abagyan, R. Methods of protein structure comparison. Methods Mol. Biol. 857, 231–257 (2012).

46 Moult, J., Fidelis, K., Kryshtafovych, A., Schwede, T. & Tramontano, A. Critical assessment of methods of protein structure prediction (CASP) – round x. Proteins 82 Suppl 2, 1–6 (2014).

47 Lovell, S. C. et al. Structure validation by Cα geometry: ϕ,Ψ and Cβ deviation. Proteins 50, 437–450 (2003).

48 Sargsyan, K., Grauffel, C. & Lim, C. How molecular size impacts RMSD applications in molecular dynamics simulations. J. Chem. Theory Comput. 13, 1518–1524 (2017).

49 Carugo, O. & Pongor, S. A normalized root-mean-square distance for comparing protein three-dimensional structures. Protein Sci. 10, 1470–1473 (2001).

50 Pettersen, E. F. et al. UCSF Chimera - A visualization system for exploratory research and analysis. J. Comput. Chem. 25, 1605–1612 (2004).

