## Supplemental Materials for "DeepPSC (protein structure camera): computer vision-based protein backbone structure reconstruction from alpha carbon trace as a case study"

**Supplementary Materials**

**Supplementary Table 1**. Fixed length (Å) used in structure rebuilding process.

| | ***trans* peptide plane** | ***cis* peptide plane** |
|---|---|---|
| $\left\|\overrightarrow{C\alpha_n PC_n}\right\|$ | 1.431 | 0.803 |
| $\left\|\overrightarrow{PC_n C_n}\right\|$ | 0.528 | 1.300 |
| $\left\|\overrightarrow{C\alpha_{n+1} PN_n}\right\|$ | 1.407 | 0.815 |
| $\left\|\overrightarrow{PN_n N_n}\right\|$ | 0.390 | 1.215 |
| $\left\|\overrightarrow{C\alpha_n PO_n}\right\|$ | 1.651 | 0.227 |
| $\left\|\overrightarrow{PO_n O_n}\right\|$ | 1.739 | 2.376 |
| $TC\beta$ | 0.914 | |
| $\left\|\overrightarrow{C\alpha_n C\beta_n}\right\|$ | 1.535 | |

**Supplementary Figure 1**. View orientation for Protein Structure Camera (PSC). This figure is generated with the Chimera software (*1*).

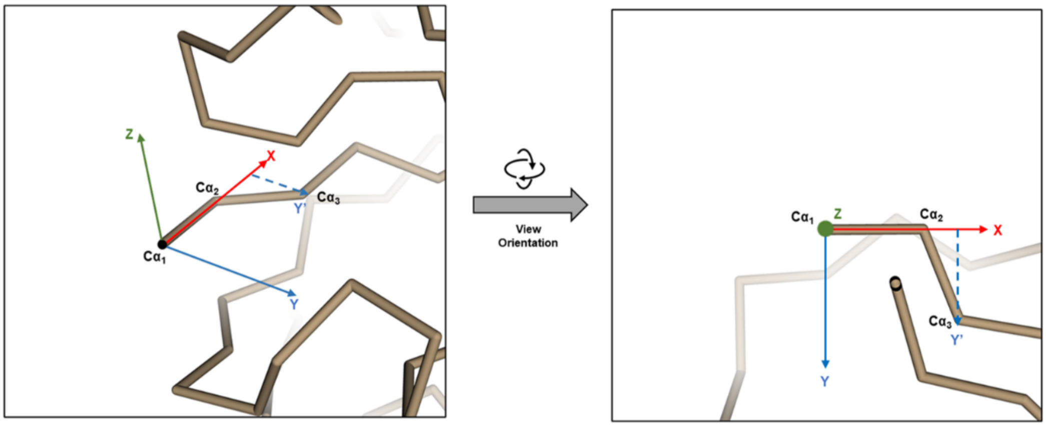

**Supplementary Figure 2**. Architecture of local structure embedding block in DeepPSC. Blocks in figure represent operations. Arrows represent data flow. The shapes of data are denoted beside arrows.

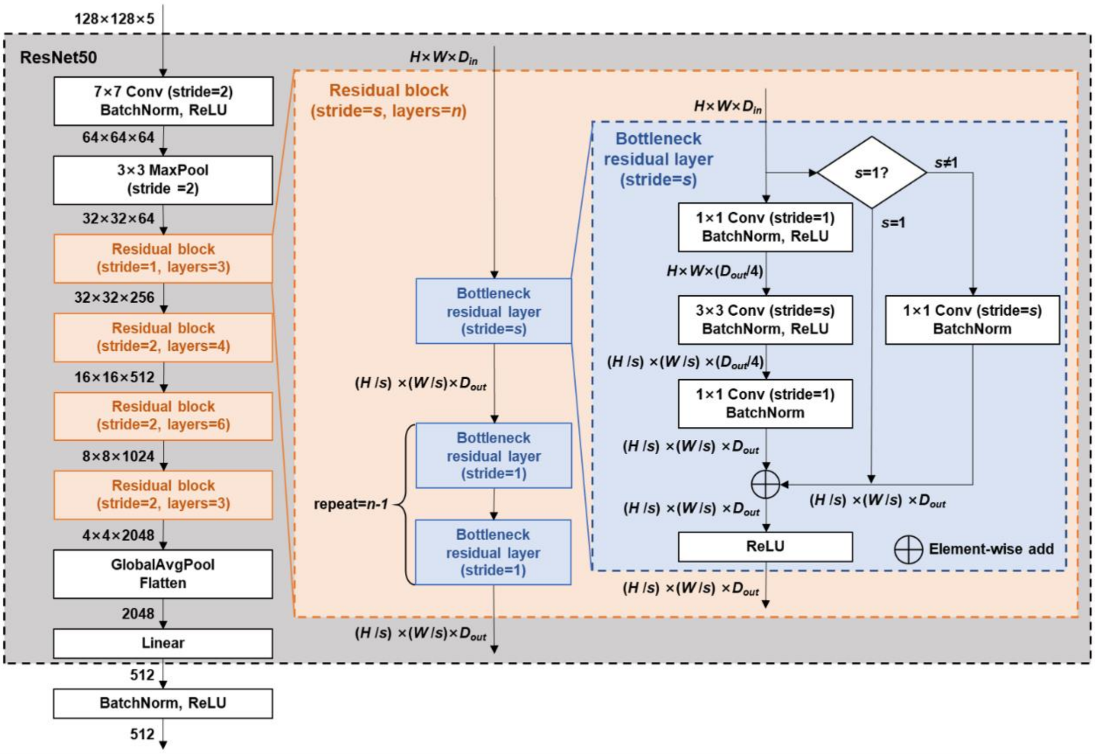

**Supplementary Figure 3**. Architecture of (a) local structure feature globalizing block and (b) prediction block in this study. Blocks in figure represent operations. Arrows represent data flow. The shapes of data are denoted beside arrows. * For baseline 2, the input dimension of local structure feature globalizing block is 32, fitting the output dimension of its local structure embedding block.

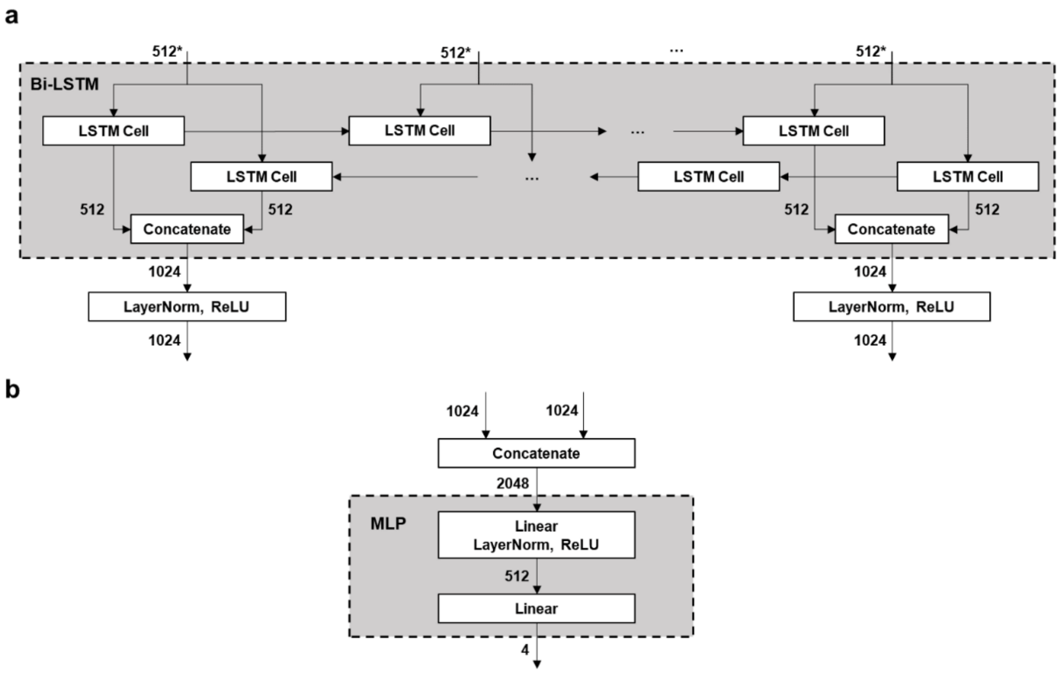

**Supplementary Figure 4**. Architecture of local structure embedding block in Baseline 1. Blocks in figure represent operations. Arrows represent data flow. The shapes of data are denoted beside arrows.

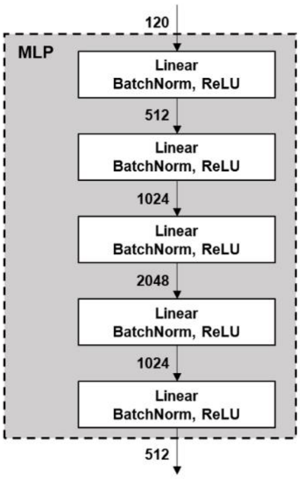

**Supplementary Figure 5**. Architecture of local structure embedding block in Baseline 2. Blocks in figure represent operations. Arrows represent data flow. The shapes of data are denoted beside arrows.

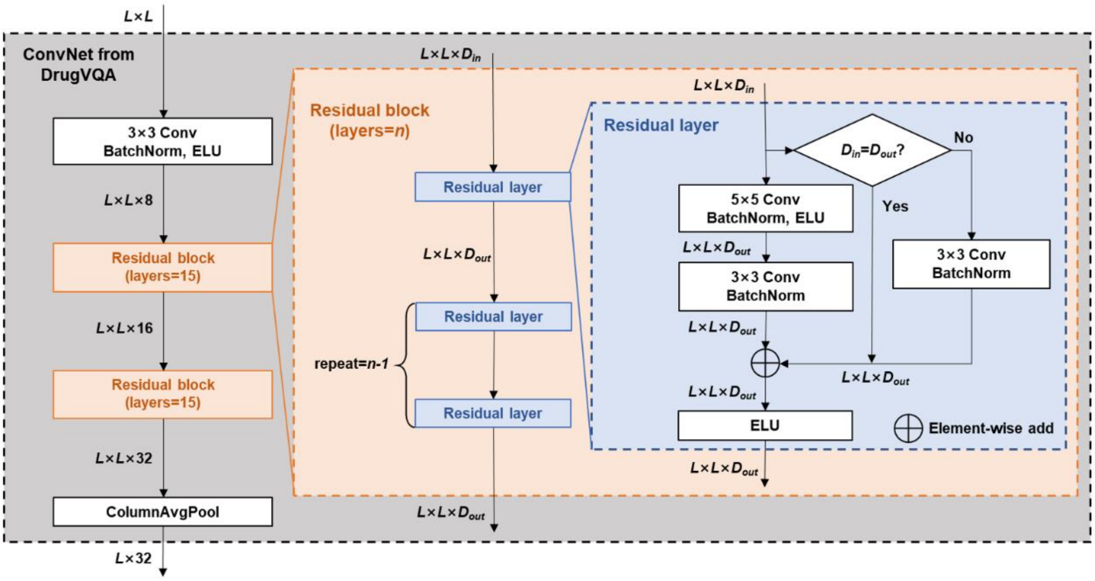

**Supplementary Figure 6**. Validation loss curves of DeepPSC, and baselines 1 and 2. For DeepPSC and baseline 1, the losses steadily decreased during the first three epochs and reached a minimum for every two epochs since the 4th epoch, due to cosine learning rate adjusting, and finally reached a minimum of 0.014 for DeepPSC, and 0.030 for baseline 1, whereas the loss for baseline 2 fluctuated around 0.183.

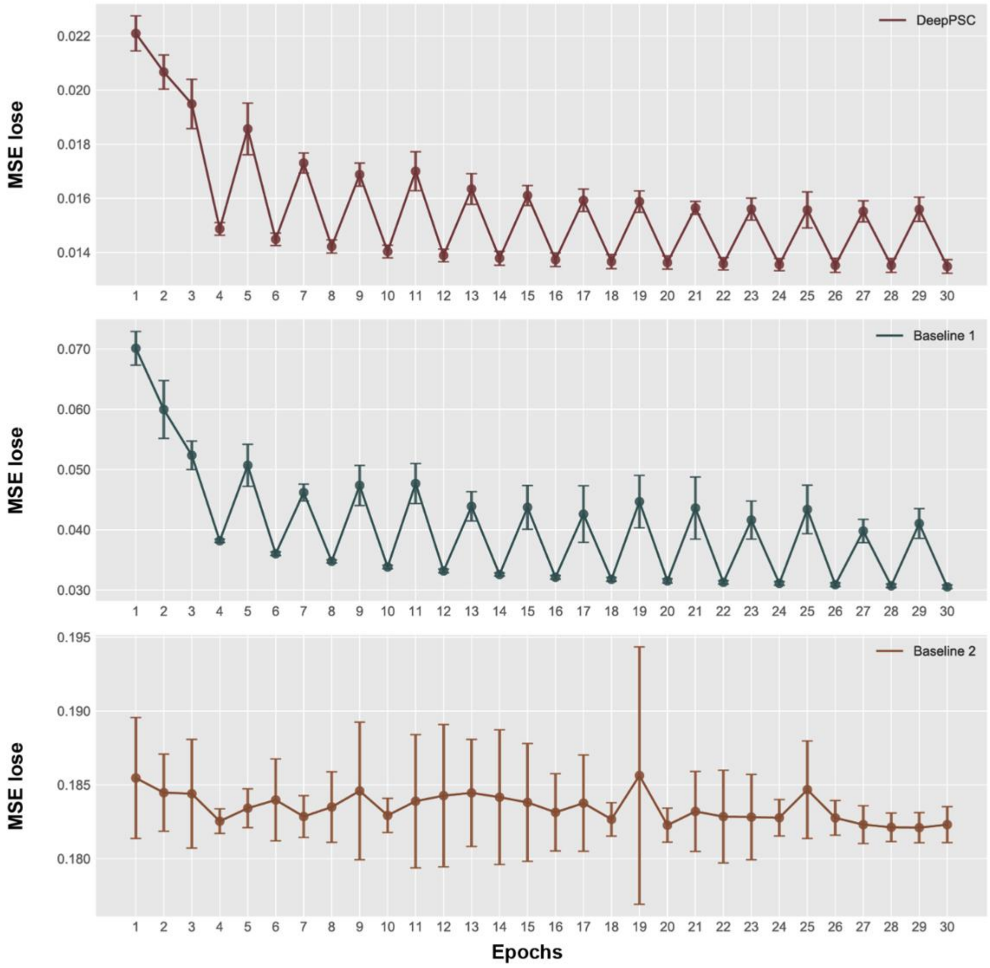


### References


1. E. F. Pettersen *et al.*, UCSF Chimera - A visualization system for exploratory research and analysis. *J. Comput. Chem.* **25**, 1605-1612 (2004).
