## Supplementary figures and images for "DeepPSC (protein structure camera): computer vision-based protein backbone structure reconstruction from alpha carbon trace as a case study"

### Supplementary Figure 1

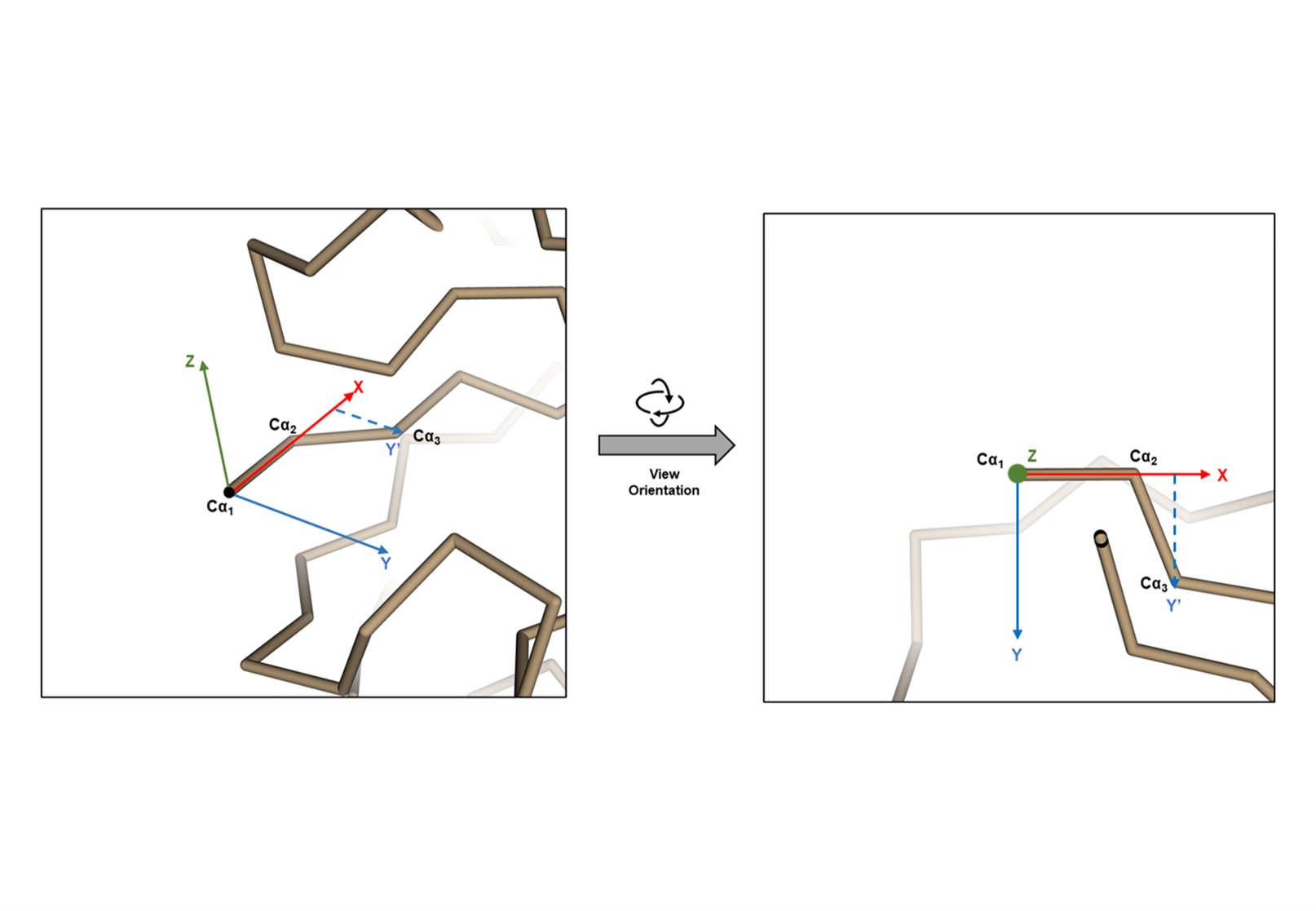

### Supplementary Figure 2

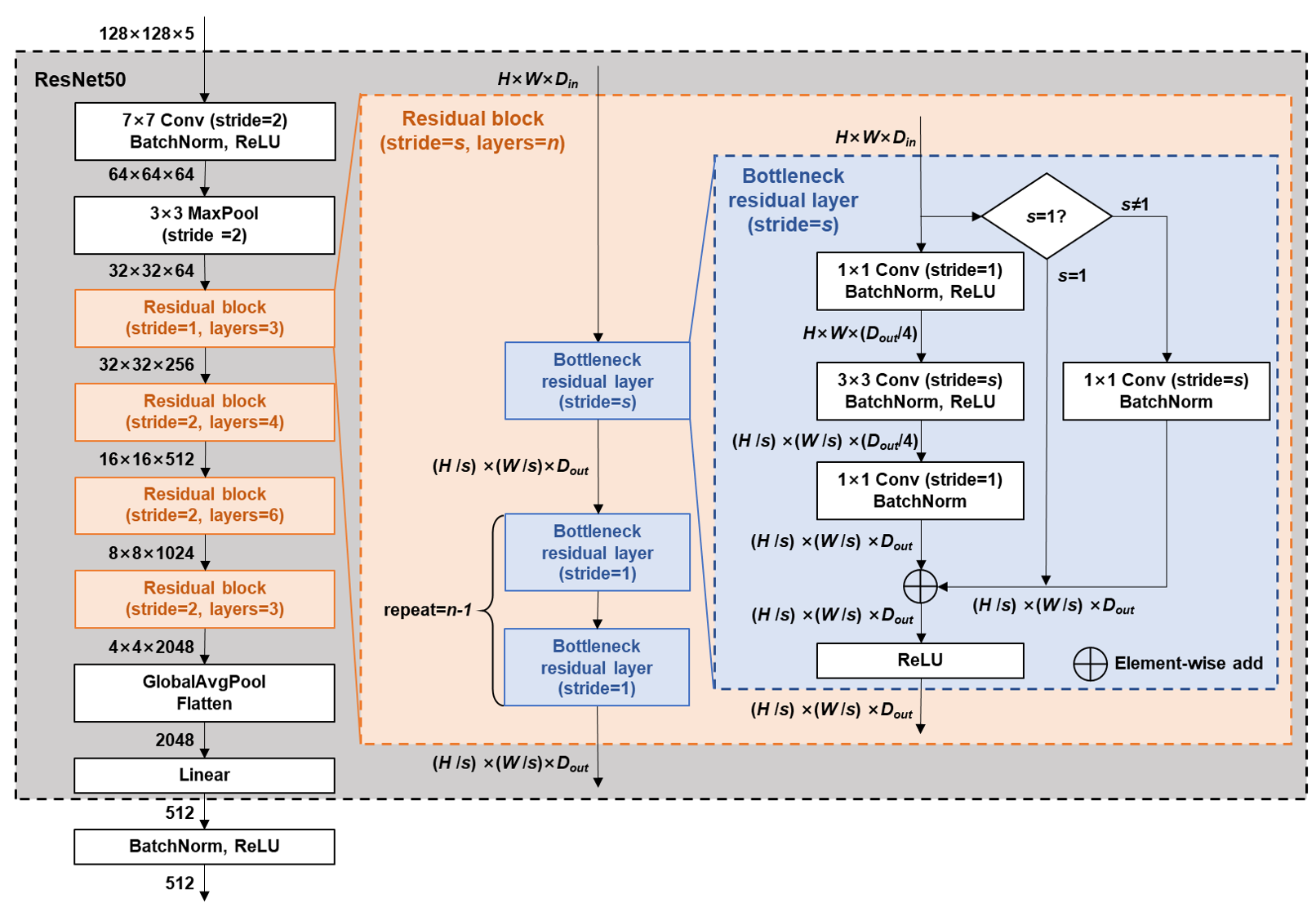

### Supplementary Figure 3

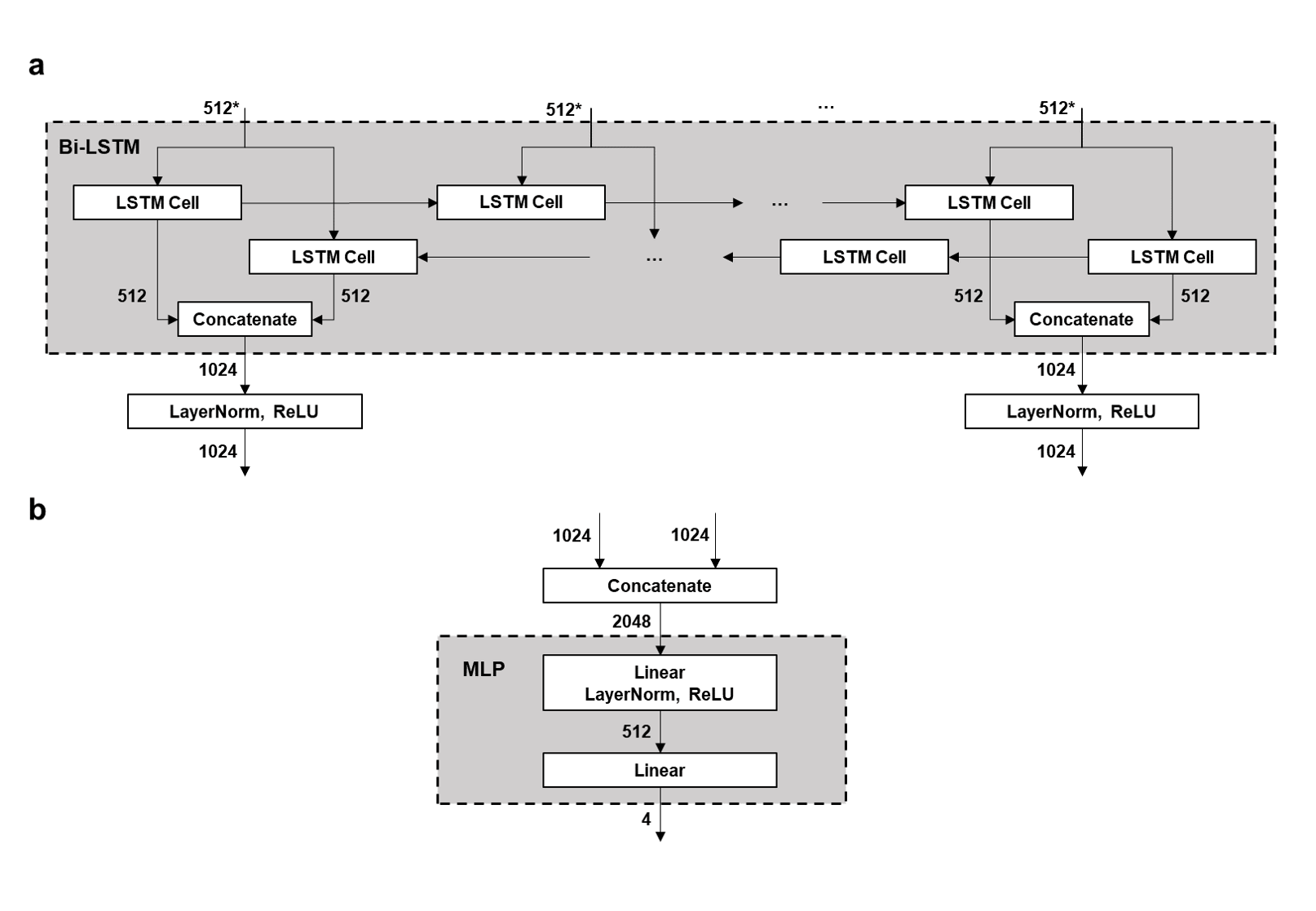

### Supplementary Figure 4

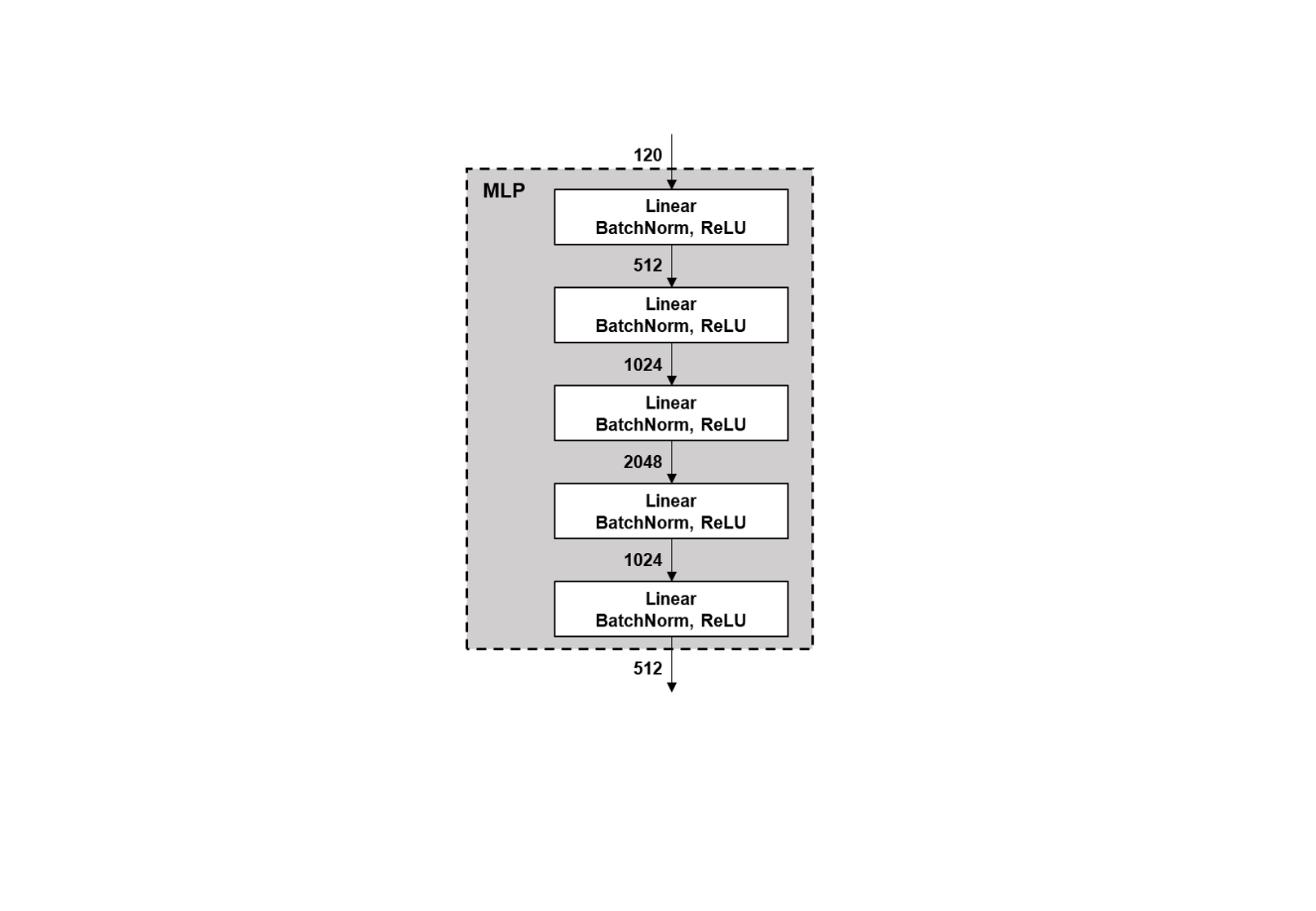

### Supplementary Figure 5

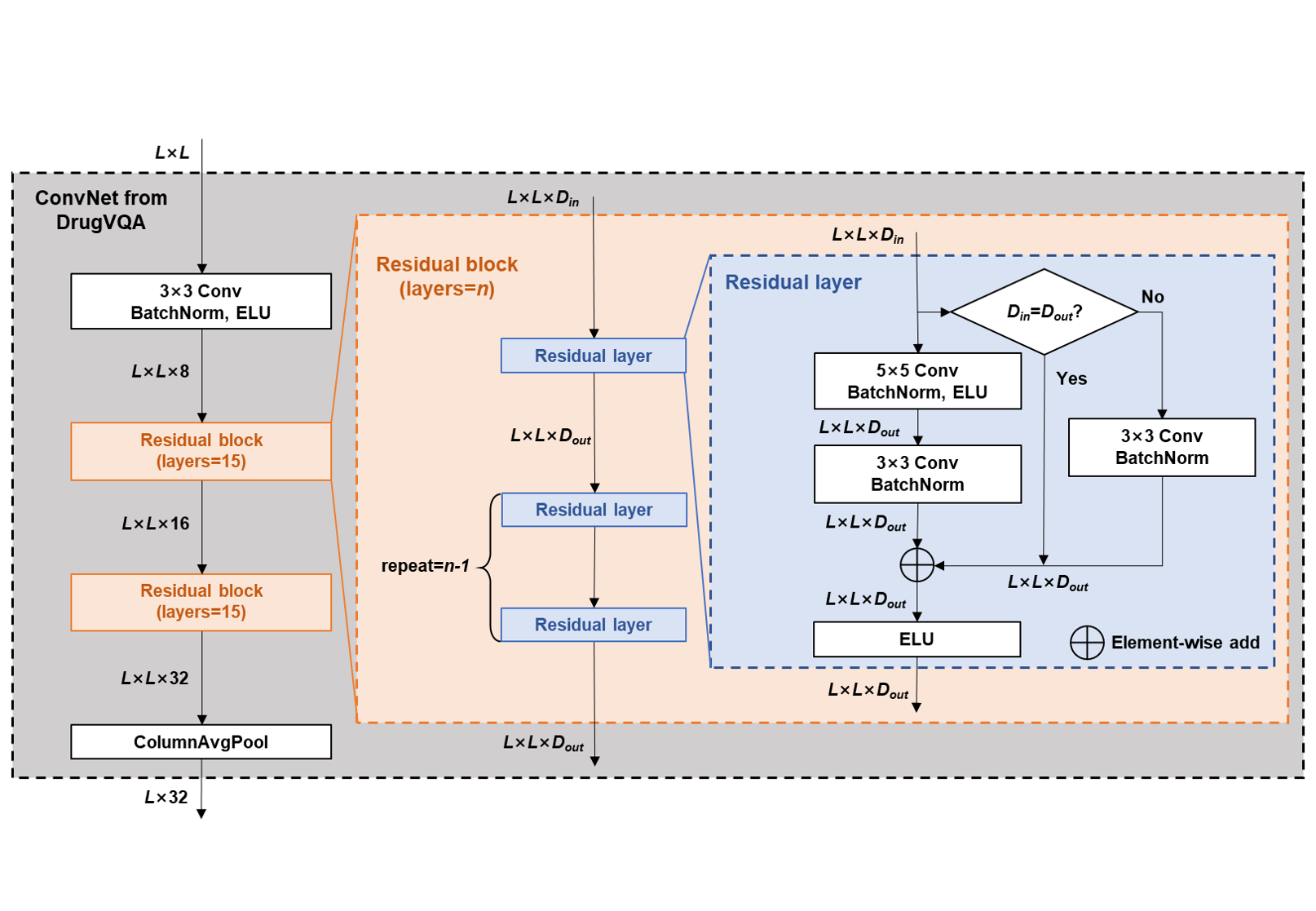

### Supplementary Figure 6

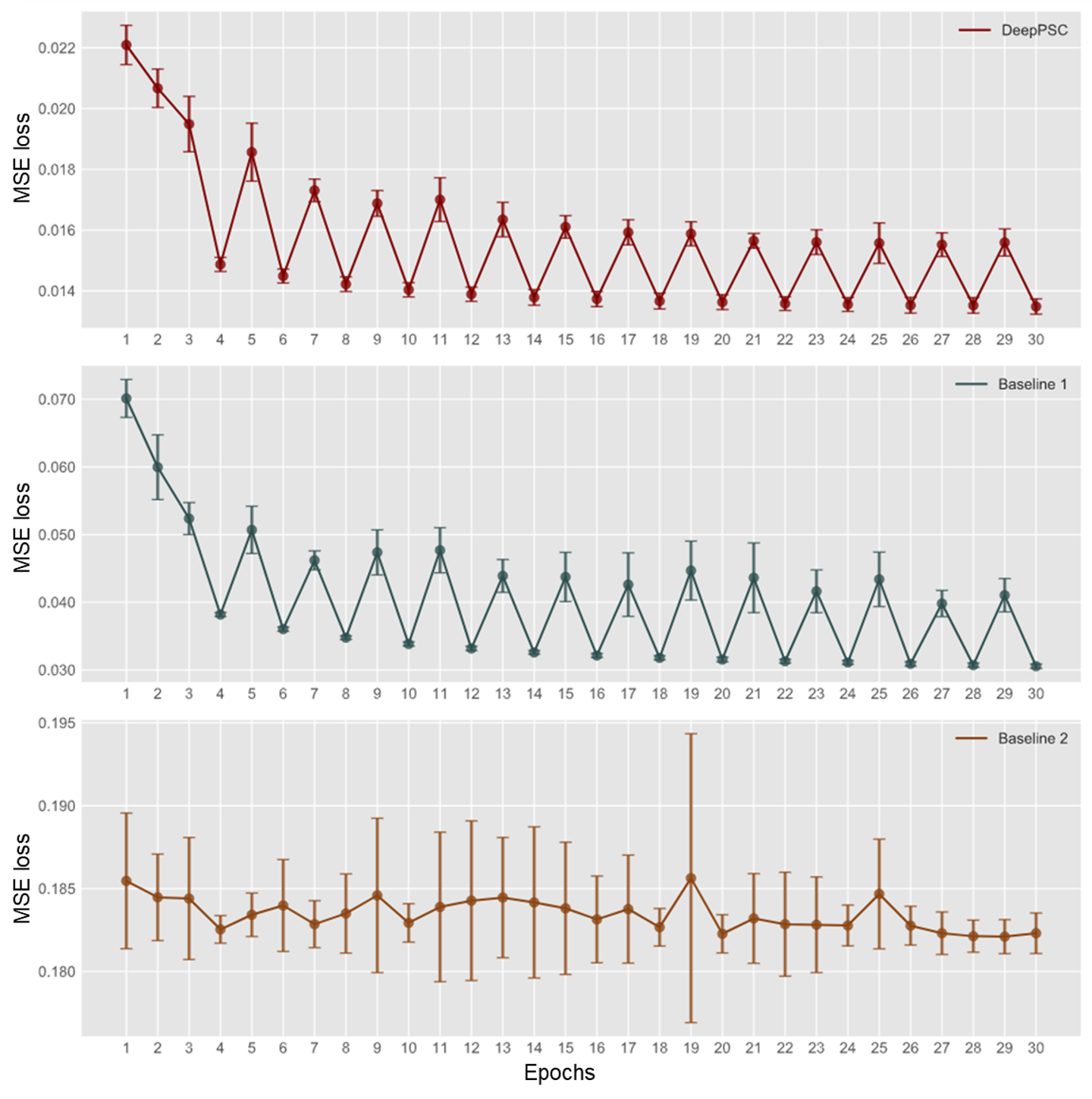
